# Drivers of large-scale spatial demographic variation in a perennial plant

**DOI:** 10.1101/2020.02.29.969428

**Authors:** Gesa Römer, Ditte M. Christiansen, Hendrik de Buhr, Kristoffer Hylander, Owen R. Jones, Sonia Merinero, Kasper Reitzel, Johan Ehrlén, Johan P. Dahlgren

## Abstract

To understand how the environment drives spatial variation in population dynamics, we need to assess the effects of a large number of potential drivers on the vital rates (survival, growth and reproduction), and explore these relationships over large geographical areas and long environmental gradients. In this study, we examined the effects of a broad variety of abiotic and biotic environmental factors, including intraspecific density, on the demography of the forest understory herb *Actaea spicata* between 2017 and 2019 at 40 sites across Sweden, including the northern range margin of its distribution. We assessed the effect of potential environmental drivers on vital rates using generalized linear mixed models (GLMMs), and then quantified the impact of each important driver on population growth rate (λ) using integral projection models (IPMs). Population dynamics of *A. spicata* were mostly driven by environmental factors affecting survival and growth, such as air humidity, soil depth and forest tree species composition, and thus those drivers jointly determined the realized niche of the species. Soil pH had a strong effect on the flowering probability, while the effect on population growth rate was relatively small. In addition to identifying specific drivers for *A. spicata’s* population dynamics, our study illustrates the impact that spatial variation in environmental conditions can have on λ. Assessing the effects of a broad range of potential drivers, as done in this study, is important not only to quantify the relative importance of different drivers for population dynamics but also to understand species distributions and abundance patterns.

## Introduction

Abiotic and biotic environmental conditions such as climate and competition vary across time and space and drive the population dynamics of species through effects on vital rates such as survival, growth and reproduction (Bruna and Oli 2005, Doak and Morris 2010). These vital rates vary both with regards to how sensitive they are to different environmental factors and to how much they influence population dynamics (Silvertown et al. 1993, Pfister 1998, Nicolè et al. 2011, Ehrlén and Morris 2015). Yet, underlying demographic mechanisms are not accounted for in standard models of species distributions and abundances (Guisan and Thuiller 2005, Araújo and Rahbek 2006, Elith and Leathwick 2009; but see Merow et al. 2017). Moreover, species distribution models based on occurrence patterns are typically based on the assumption that species are in equilibrium with their environment, and thus do not capture ongoing or recent changes in environmental drivers. To achieve an in-depth understanding of the population dynamics of a species, and to describe and predict distributions and abundances, we need to assess how environmental variation is linked to demographic variation (Ehrlén and Morris 2015). Using environmentally explicit demographic models allows us to disentangle which vital rates are most affected by the environment, and how those vital rates contribute to population growth rate *(λ).* Such a more mechanistic understanding of the processes underlying a species’ distribution and abundance allows for more accurate predictions of species responses to environmental and climate changes (Doak and Morris 2010, Csergő et al. 2017). Environmentally explicit demographic models also provide a means to identify the drivers of the short-term dynamics and of the realized niches of species.

To avoid preconceived notions about which drivers are important, assessments of environmental effects on population dynamics should include a broad range of potential drivers, ideally including climatic and other abiotic factors, as well as biotic and anthropogenic factors such as land use (Ehrlén et al. 2016). In addition to assessing many potential drivers, data collection covering as big a part of the environmental gradients a species experiences as possible should allow for more reliable models (Ehrlén et al. 2016). Meeting these needs regarding the number of drivers and accurately estimating their variation will likely often mean that data have to be collected over large geographical areas. However, although the modelling tools for assessing the effects of environmental conditions on population dynamics and distributions are available (Merow et al. 2014*a*, 2014*b*), few studies have investigated the relationships between multiple vital rates and environmental drivers on large geographical scales (but see Merow et al. 2017).

We conducted a large-scale study of the environment-driven demography of the long-lived understory forest herb *Actaea spicata* based on data from a large geographical extent. We collected demographic data as well as data on 18 putative environmental drivers, from 40 populations throughout the Swedish distributional range of *A. spicata* from 2017 to 2019. The putative drivers were 15 abiotic factors related to climate, topography and soil nutrient richness, and three biotic factors concerning plant community structure and intraspecific population density. We asked the following questions: (1) How do the various environmental factors influence the vital rates of *A. spicata*? We expected strong effects of both the physical environment, in terms of nutrient levels, and the biotic environment, in terms of shading by other plants, in line with previous smaller-scale studies (Dahlgren and Ehrlén 2009, 2011). Specifically, we expected soil potassium concentrations to positively influence growth and increasing proportion of coniferous trees to negatively influence growth (Dahlgren and Ehrlén 2009, 2011). Furthermore, we expected strong climatic effects due to the large geographical range of the study, and the fact that the study was conducted at the northern range limit of *A. spicata*. (2) How do detected effects of environmental drivers on vital rates influence the population growth rate of *A. spicata*? We expected that the environmental drivers affecting survival and growth would have a larger effect on population growth rate than those affecting reproduction, because such dynamics are described for other long-lived herbs (Silvertown et al. 1993). We used generalized linear mixed models (GLMM) to link the environmental drivers to vital rates describing survival, individual growth, flowering probability and fruit number. We then incorporated the GLMMs into an integral projection model (IPM, Easterling et al. 2000) to assess the effects of environmental drivers on the population growth rate of *A. spicata.*

## Methods

### Study species

Baneberry *(Actaea spicata* L., Ranunculaceae) is distributed over most of Europe and parts of Asia and North America (Anderberg A. and Anderberg A.-L. 2017). In northern Europe, it occurs in shady, well-drained and nutrient rich forests, often on limestone (Pellmyr 1984). The plant is common throughout Sweden but does not occur in the far north (Mossberg and Stenberg 2014). *Actaea spicata* has a morphology typical of many early summer flowering forest herbs, with a greater height than spring ephemerals and an umbrella-like leaf display (cf. Givnish 1987). Previous studies have found that individuals can produce several shoots, each typically bearing up to four inflorescences (Eriksson 1995), with the first, top inflorescence, typically bearing most flowers and fruits. Each black berry contains 8-16 seeds (Zeipel et al. 2006) which germinate below ground one year after release, and aerial parts emerge one year thereafter (Ehrlén and Eriksson 2000, Fröborg and Eriksson 2003). The entire plant is toxic (Anderberg A. and Anderberg A.-L. 2017). Estimated mean life span of individuals surviving to reproduction is 20.2 years (Ehrlén and Dahlgren, unpublished data). Population growth rate is more sensitive to survival and growth than to reproduction (Dahlgren and Ehrlén 2009), as is typical for long-lived herbs (Silvertown et al. 1993).

### Study area

Potential study populations were selected using Artportalen (www.artportalen.se), an online reporting system for species observations (including citizen science observations) and information about Swedish flora and fauna developed by the Swedish University of Agricultural Sciences (SLU). We visited more than 100 populations across the country distributed with the aim of covering the entire Swedish distribution of the species and to obtain a representative sample of the environments experienced by the species (favoring reported occurrences with more exact coordinates). We aimed for populations with at least 50 established non-seedling individuals, but occasionally populations with fewer individuals were included in the study to guarantee a good spatial coverage. We identified 43 suitable populations across Sweden (Appendix S1: Fig. S1). In these populations permanent plots were established in 2017 and followed until 2019. Over the course of the study, we lost three populations due to grazing and wild boar activity, leaving 40 study populations in which data was collected in all three years. Distance to nearest population ranged between 30 km and 139 km (mean distance = 73 km) and plot size depended on the plant density and ranged from 6 to 363 m^2^. The habitat varied substantially between sites, with some sites located in pure broadleaf forest and others in pure coniferous forests.

### Demographic data collection

Up to 50 non-seedling individuals were included at each population in 2017. All plants were marked by placing a small flag (steel wire with a piece of tape) in the soil, and mapped to enable relocation in following years. All seedlings were counted and followed individually from the year after emergence. In total, 2263 plants were surveyed over the course of the study. Presence (1 or 0), state (flowering or vegetative), plant height, shoot diameter, and fruit number per inflorescence were recorded for all established plants between July and August in all years.

An individual was considered dead in 2018 if it lacked above-ground structures in both 2018 and 2019. Individuals that were only missing in 2018 had either been dormant or the above-ground tissues had been damaged before the demographic census. Because dormancy in one year could only be detected in retrospect after revisiting the site in the following year, we assumed the same proportion (3.6%) of individuals lacking above-ground structures were dormant in 2019 in the statistical models for survival. Plant size was defined as the natural logarithm of the product of plant height and squared shoot diameter, where height is the distance from the ground to the horizontal plane formed by the largest leaves (as in Dahlgren and Ehrlén 2009; see Appendix S2: section 1). Growth was defined as the difference in size between censuses.

### Environmental data collection

We collected a range of environmental variables including 15 abiotic factors related to climate, topography and soil properties, and three biotic factors concerning plant community structure and intraspecific population density. We collected weather data throughout the whole study period, with air-temperature and humidity data loggers (EasyLog EL-USB-2, Lascar electronics) placed at the center of each plot and covered with a plastic cup to protect them from direct sunlight and rain. We used the logger data to calculate (i) growing degree days (base temperature: 5 °C) for the spring period 15th May-15th June.06, (ii) the start of spring, using the definition from the Swedish meteorological office (SMHI 2011) and (iii) vapor pressure deficit (hereafter VPD), the difference between the actual vapor pressure in the air and that of saturated air, where vapor pressure is a temperature corrected measure for moisture (Anderson 1936, Jones 2014). VPD was calculated for the summer period (15th June – 15th August) using the R package *plantecophys* (Duursma 2015). We measured slope inclination and slope aspect and used them to calculate the extent to which the ground was slanted towards the sun, as an additional indicator of the exposure to light and moisture (hereafter “slope”). In addition, we recorded exact latitude and longitude positions with a Garmin GPS map 64s, estimated site area and calculated intraspecific plant density for each site as the number of individuals per m^2^. In 2018 we collected soil samples and analyzed the concentrations of nitrate (NO_3_^-^), exchangeable ammonia (NH_4_^+^), plant available phosphorus (P), phosphate (PO_4_^3-^), plant available potassium (K) as well as soil pH. Lastly, we collected data on the plant community structure at each site: To determine the percentage of coniferous vs. broadleaf trees we recorded the abundance of each tree species using a relascope, a forestry instrument to measure basal area, for each tree visible from the center of the site. We took canopy cover pictures using the back camera of a Sony Xperia L1 with an attached fisheye-lens (180° Supreme Fisheye Lens, Model MFE4 by MPOW Inc) which were then processed in ImageJ (Schneider et al. 2012) using the plugin Hemispherical 2.0 (Beckschäfer 2015) to calculate the gap fraction (see Appendix S2: section 2 for further details on the environmental data collection).

**Table 1:**
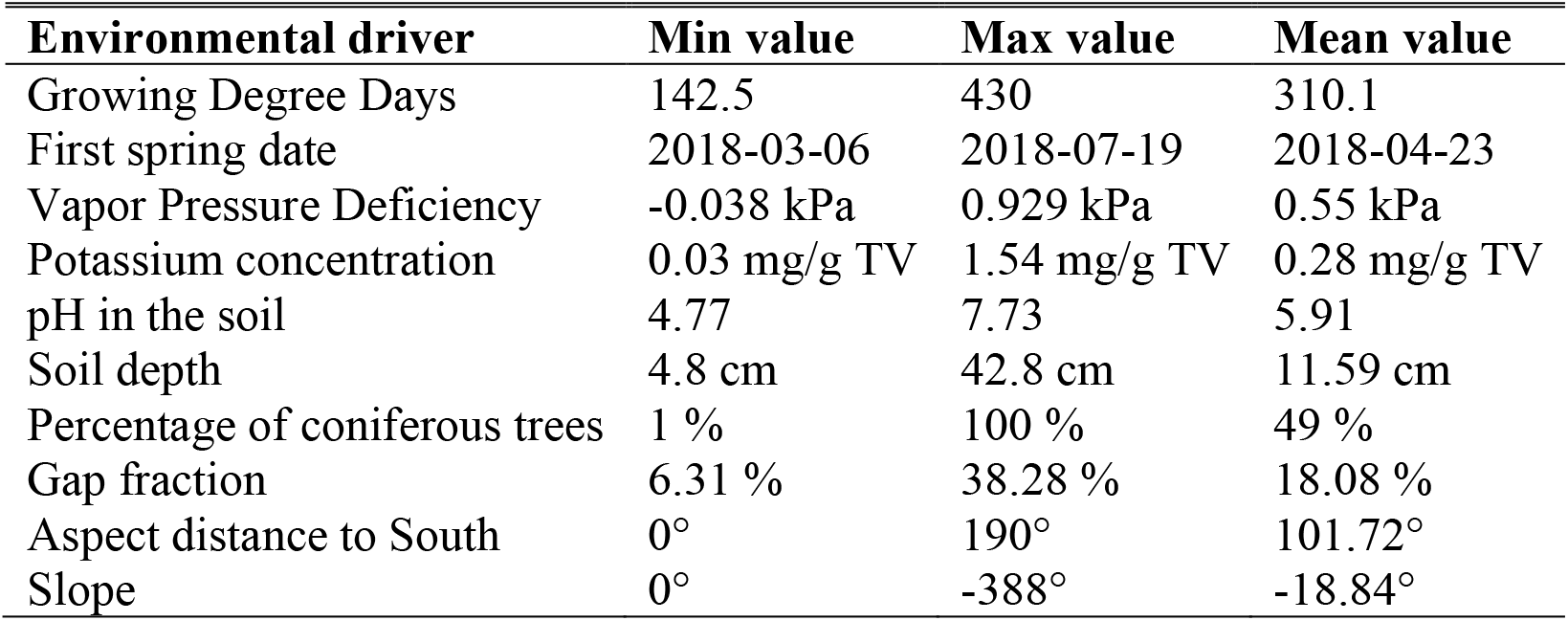
Observed values of environmental drivers in the field. For each environmental driver we present minimum and maximum observed value on our 40 field sites as well as the mean of all observations.

### Statistical analyses

To avoid the effects of overfitting (Harrell 2001), we limited the number of environmental variables included in the regression models, and did not evaluate interactive effects. If two environmental variables showed bivariate correlations of > 0.7, we omitted the variable we expected less likely to have an effect on the population growth rate of *A. spicata* based on knowledge of the system (Appendix S1: Fig. S2). To further reduce the number of putative drivers, we performed a principal component analysis (PCA) on the soil variables. All soil variables except pH were aligned with the first PCA-axis, which explained 46.8% of the variation, and soil pH was aligned with the second PCA-axis, which explained 22.3% of the variation (Appendix S1: Fig. S3). Based on this, we chose to retain only soil potassium and pH as soil potassium has previously been shown to have important effects on *A. spicata* (Dahlgren and Ehrlén 2009, 2011). Final models thus included the following environmental variables in the vital rate regression models: VPD, soil potassium concentration, soil pH, soil depth, percentage of coniferous trees, intraspecific density, canopy gap fraction and slope.

We assessed effects of environmental variables on vital rates (probability of survival, growth, probability of flowering, and number of fruits) using generalized linear mixed models (GLMM) and linear mixed models (LMM) in the R package *lme4* (Bates et al. 2015). Predictions lines and 95% confidence intervals were generated with the R package *effects* (Fox and Weisberg 2018). We standardized all variables by subtracting the arithmetic mean and dividing by the standard deviation to ease interpretation when comparing effect sizes and to improve model convergence. For modelling the probability of survival and flowering we used logistic regressions with binomial error distributions and logit link functions. Growth was modelled with a ordinary Gaussian linear model and fruit number was modelled using a Poisson error distribution and log link function. We pooled the data for both annual transitions for analysis and accounted for the spatial structure of the data and repeated measurements of individuals by including site and plant ID as random effects. We also analyzed the two years separately, to investigate the consistency of effects. In all models, we included plant size as a fixed effect.

We included all environmental variables and intraspecific population density as both linear and quadratic fixed-effect terms. To ease interpretation and avoid effects of overfitting (Harrell 2001) we then created a reduced model, which only included those quadratic terms that where statistically significant following strict cut-off points (for the GLMMs: *P* < 0.05; for the LMMs: no *P*-values are provided by the *lme4* package, we therefore used – 1.96 < *t* < 1.96 which corresponds to a *P*-value of < 0.05 in models with high degrees of freedom). We used this reduced model to assess the statistical significance of remaining linear and quadratic terms, and we evaluated the statistical significant of effects according to the same criteria as above. We conducted all modelling, data exploration and visualization using R version 3.5.1 (R Core Team 2018).

### Integral Projection Modelling

An IPM is a population model with similar properties as a matrix population model but with continuous instead of discrete stage classes and a projection kernel instead of a projection matrix (Easterling et al. 2000). We constructed the IPM in this study as described in Dahlgren and Ehrlén (2009) based on R code presented in Merow et al. (2014*a*, Appendix F). To allow for a delay of seedling establishment (see Study species) and treatment of seeds and seedlings as discrete classes (cf. Rees et al. 2006) our environmentally explicit IPM consisted of three coupled equations:

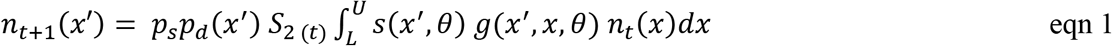

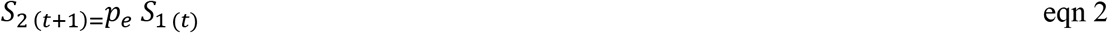

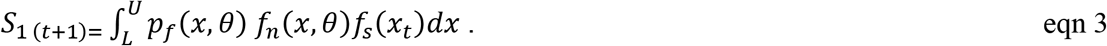

Each relevant component of the IPM included a vector *θ* for the significant environmental drivers identified in the regression models as covariates. *n(x_t_)dx* is the distribution of size *x* at time *t* while *L* and *U* are the upper and lower bounds of possible sizes. In equation 1, *p_s_* is the probability of seedling survival and *p_d_* the probability density function of sizes of surviving seedlings. S_1_ is the number of seeds and S_2_ the number of seedlings. The function *s(x′, θ)* describes survival and *g(x′, x, θ)* is the growth function, describing individuals of size *x* at time *t* which survive reaching size *x′* in time *t+1.* In equation 2, *p_e_* is the probability of seedling establishment given the survival of seeds (*S_1_*). In equation 3,*p_f_*(*x*) is the probability of flowering and *f_n_* the number of fruits. *f_s_* is the number of seeds per fruit, which was assumed to be 9.61, as in Dahlgren and Ehrlén (2009).

We first used the IPM to calculate population growth rate *(λ)* as the dominant eigenvalue of the kernel (Ellner and Rees 2006, Ellner et al. 2016) for each study site. We then estimated the correlation between *λ* and latitude using a linear regression model. To evaluate the effects of each significant environmental variable on *λ,* we constructed a separate kernel for each environmental variable and calculated *λ* for the observed mean values of the tested variable as well as for a range corresponding to three standard deviations away from the mean (i.e. ranging from −3 to 3) while holding all other variables constant at their mean value.

## Results

Over the two observed transitions the average yearly survival of individuals was 91%. The mean change in size from year *t* to year *t*+1 was −3.6%. On average 24% of all individuals flowered and the number of fruits per individual ranged from 0 to 90 (mean = 16.15). The first inflorescence carried on average 82% of all fruits.

Individual plant size was correlated with all four investigated vital rates; survival, growth, probability of flowering, and number of fruits (Appendix S1: Fig. S4 and table S1). Soil depth, soil pH, VPD, intraspecific density and percentage of coniferous trees were significantly related to one or several vital rates (Fig. 2). Most environment–vital rate relationships were linear, but density and percentage of coniferous trees had significant quadratic relationships with fruit number (Fig. 2d). In our regression models, the percentage of coniferous trees affected survival positively (Fig. 2a) whereas fruit number was highest at average values of percentage of coniferous trees (Fig. 2d). VPD negatively influenced growth and flowering probability (Fig. 2b and c). Soil depth had a positive relationship on growth (Fig. 2b). Soil pH was negatively related to flowering probability (Fig. 2c) and the effect of intraspecific density on number of fruits was u-shaped with higher fruit production at lowest and highest density values (Fig. 2d). The effects of environmental variables were similar in both transitions (Appendix S1: table S4).

**Figure 1.**
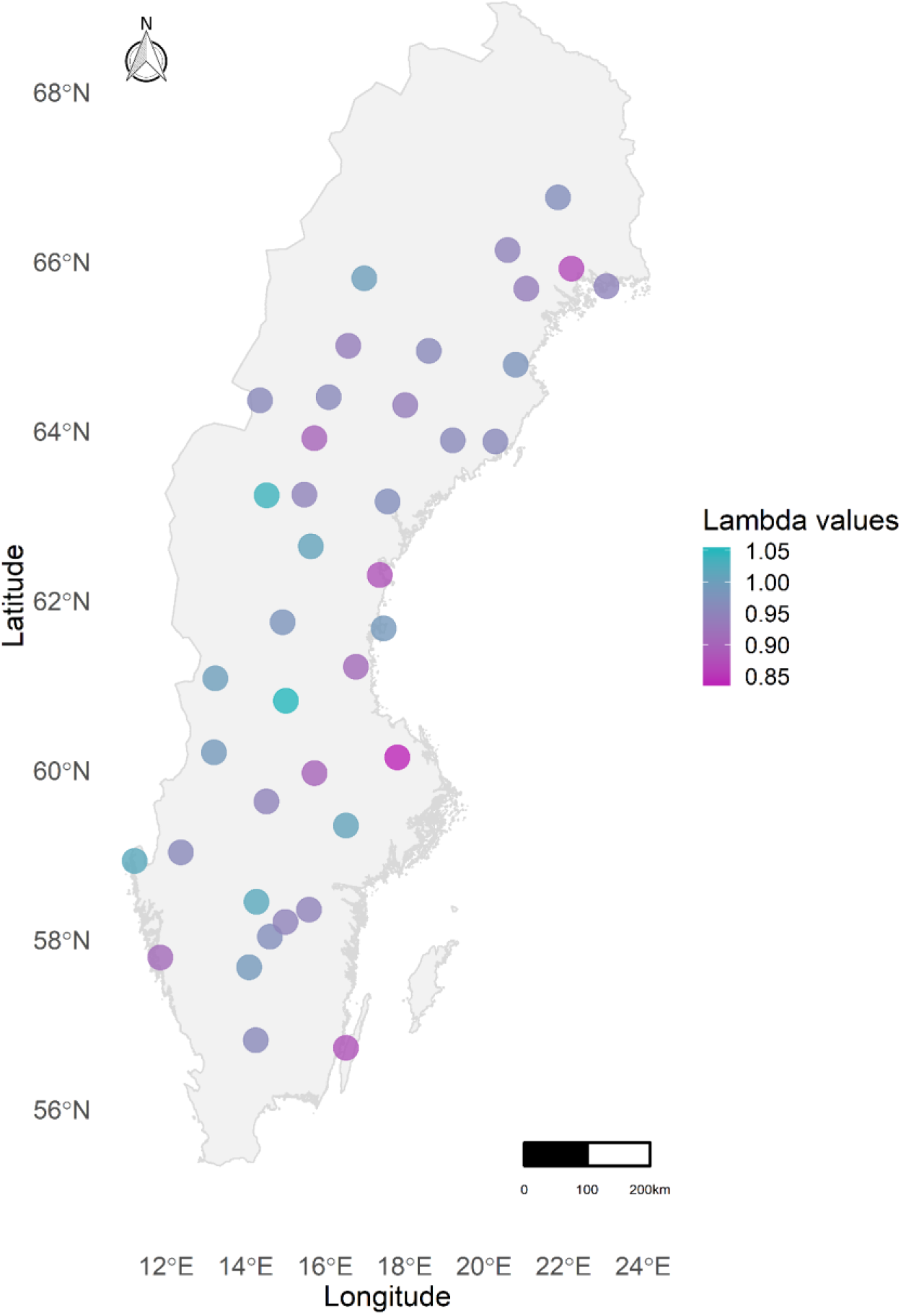
Map of Sweden showing the population growth value (*λ*) for the corresponding site for the time period 2017-2019.

**Figure 2.**
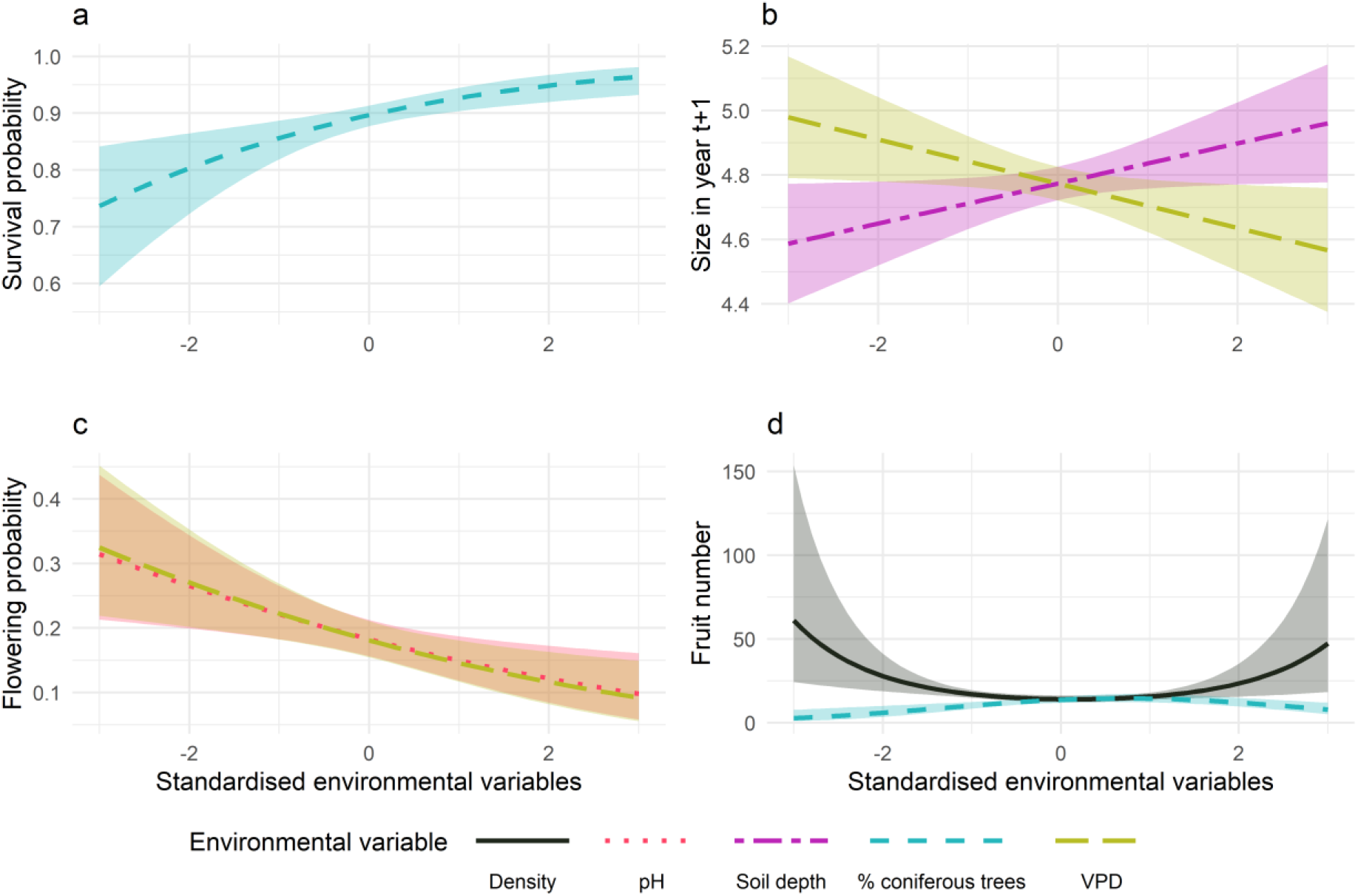
Influence of environmental drivers on vital rates of *A. spicata*. Panel (a) shows the influence of coniferous trees (β = 0.38, ± = 0.11 SEM, *P* = < 0.01) on survival, (b) VPD (β = −0.07, ± = 0.03 SEM, *t* = −2.209) and soil depth (β = 0.06, ± = 0.03 SEM, *t* = 2.071) on growth, (c) pH (β = – 0.24, ± = 0.09 SEM, *P* = < 0.01) and VPD (β = −0.26, ± = 0.09 SEM, *P* = < 0.01) on the probability of flowering and (d) density (β = 0.17, ± = 0.07 SEM, *P* = 0.01; β = −0.12, ± = 0.04 SEM, *P* = < 0.01 for density and density^2^, respectively) and percentage of coniferous trees (β = −0.04, ± = 0.04 SEM, *P* = 0.25; β = 0.15, ± = 0.06 SEM, *P* = < 0.01, for coniferous trees and coniferous trees^2^, respectively) on the number of fruits. Polygons show 95% confidence intervals.

The mean population growth rate was 0.96 over all transitions and sites. However, populations were on average increasing during the first transition (mean λ_2017-2018_ = 1.04, sd = 0.05), but decreasing during the second transition (mean λ_2018-2019_ = 0.87, sd = 0.05). Population growth rates also varied between sites (Fig. 1) and ranged from 0.84 to 1.05 over the study period (sd = 0.05, Appendix S1: table S2). We found no significant relationship between population growth rate and latitude *(P* = 0.77).

Our IPMs showed that coniferous trees and VPD had the largest influence on population growth rate, closely followed by the effect of soil depth (Fig. 3, Appendix S1: Fig. S5 and table S3). While the effects of increasing proportion of coniferous trees and soil depth were positive, increasing VPD had a negative association with *λ*. The influence of increasing pH was also negative but smaller than the effect of VPD. The effect of density was negligible.

**Figure 3.**
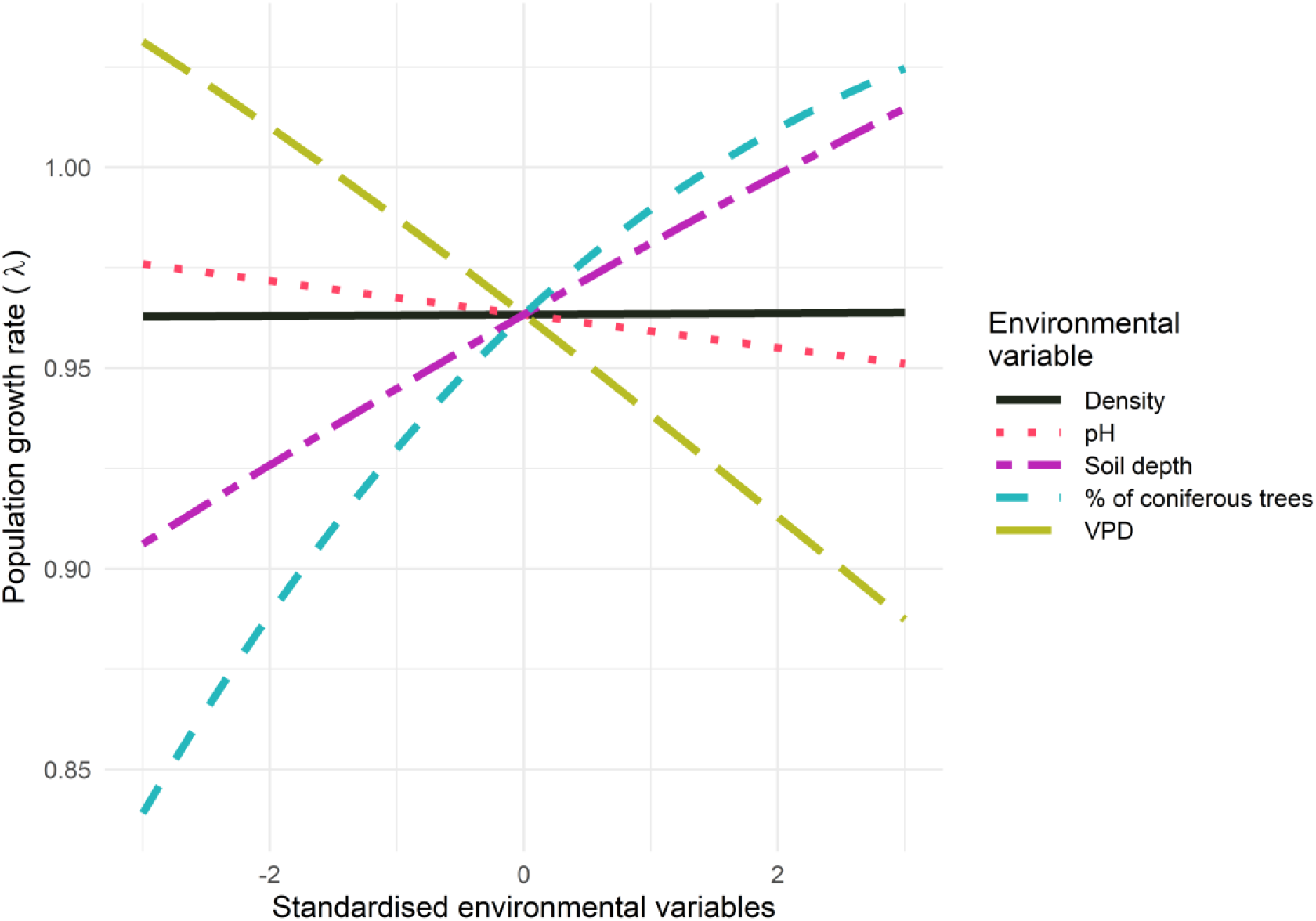
Influence of the environmental drivers of vital rates on the population growth rate (*λ*).

## Discussion

Our results demonstrate substantial effects of climatic, non-climatic abiotic, and biotic environmental drivers on the population dynamics of the perennial herb *Actaea spicata*. Specifically, we showed that population growth rate is strongly influenced by vapor pressure deficit (VPD), soil depth, and tree composition. There was also a substantial difference in the relative importance of the various environmental drivers on the different vital rates. The three environmental factors influencing survival and growth had large effects on overall population dynamics, illustrating that the population growth rate of *A. spicata,* as well as other long-lived plants, is more sensitive to changes in those vital rates (Silvertown et al. 1993, Dahlgren and Ehrlén 2009). In accordance with the low influence of reproduction on population growth rate of *A. spicata*, soil pH had small effects on population dynamics, even though it had a strong negative effect on the probability of flowering. The effect of intraspecific density on fruit number translated to an even smaller effect on population growth rate. These results illustrate how effects of environmental factors and the sensitivity to the affected vital rates jointly determine the realized niches of species by putting limits on population growth.

The negative effects of VPD on several vital rates and population performance is particularly interesting in the context of ongoing climatic changes. If humidity in Sweden will increase at the same rate as it has during the last 50 years (Wern 2013), and temperature increases as predicted by Sjökvist et al. (2015), the predicted average population growth rate of *A. spicata* across Sweden would increase from *λ* = 0.97 to *λ* = 1.01 (see Appendix S3 for details). The positive influence of lower VPD values on population growth rate, and the expectation of a future decrease, highlight the importance of understanding the effects of climate change on plant populations. Like several previous studies in temperate areas (Nicolè et al. 2011, Sletvold et al. 2013), we found that direct effects of climate change on plant populations will likely be positive. If this is a general trend, and that most species would be affected positively, then this should mean that effective competitors may be the most successful plants in the future. In addition to direct effects on *A. spicata* and its competitors, changes in climate might affect forest community structure and soil composition (e.g. Lükewille and Wright 1997, Garten et al. 1999), which in turn affect its population dynamics. In a changing climate, a combination of these factors will lead to a shift in the geographical distribution of suitable conditions for *A. spicata.*

Some of the effects that we detected contrasted with our expectations. The positive effect of a high percentage of coniferous trees on individual survival and population growth rates is the opposite of the effect found in previous studies with the same species (Fröborg and Eriksson 2003, Dahlgren and Ehrlén 2011). It also contrasts with that *A. spicata* tends to be more common in broadleaf forests (Mossberg and Stenberg 2014). It is possible that the percentage of coniferous trees in our study was correlated with, for example, low light availability, high soil moisture or low maximum temperatures. This could explain why its effect on population growth was stronger in the 2018-2019 transition following an extremely warm and dry in 2018, suggesting the positive effect of coniferous trees on survival may have been due to a buffering of the drought. Previous studies also detected effects of soil potassium, but no effects of soil pH, soil depth and variables related to soil moisture and temperature (Dahlgren and Ehrlén 2009, 2011). The differences in detected effects are likely due to the different scale of the studies, where the previous were focused on patterns at a few sites close to each other while this study covers the Swedish distribution of the species. The difference in results highlight the importance of scale for demographic studies. Large-scale studies are likely to cover larger environmental gradients and therefore more likely to detect responses to an environmental driver (e.g. Levin 1992, Chave 2013).

We detected both biotic and abiotic drivers and several of these might alter due to anthropogenic influences, such as effects of forestry on tree species composition. Our findings are in line with other studies that have assessed effects of multiple drivers of population dynamics (Nicolè et al. 2011, Diez et al. 2014) and a recent study found that the impact of abiotic, biotic and anthropogenic factors on population growth rate are equally important and should therefore all be considered in environmentally explicit population models (Morris et al. 2020). Our results emphasize that the performance of a species is affected by multiple environmental drivers, and that we should consider as many drivers as possible, given considerations of sample size and potential overfitting, when assessing environmental drivers and predicting how environmental change will affect species. We conclude that environmentally explicit demographic models, fitted to data over large geographical scales, and including a broad variety of environmental drivers are a valuable tool for identifying species’ niches. We also assert that these analyses can be valuable for designing management strategies for species of special interest. Finally, combining these demographic models with classical species distribution models (e.g. Greiser et al. 2020) has the potential to lead to a deeper understanding of the drivers of changes in species abundances and distributions.

## Acknowledgements

We thank Mads Nedergaard Olsen, Malin Borg, Ebba Tamm and Laura Lecacheux for help with planning and conducting fieldwork, as well as Carina Kronborg Lohmann and Rikke Holm for processing the soil samples. The project was supported with a grant from the Swedish Research Council for Environment, Agricultural Sciences and Spatial Planning (FORMAS) (to J. E. and J.P.D). JE and JPD conceived the overall research plan. GR, DMC, KH, ORJ, SM, JE and JPD designed fieldwork. GR and HdB conducted field work. KR conducted soil analyses. GR analyzed data with assistance from DMC and JPD. GR, JPD and JE led the writing of the manuscript, with input from all authors.

## Supplementary Information

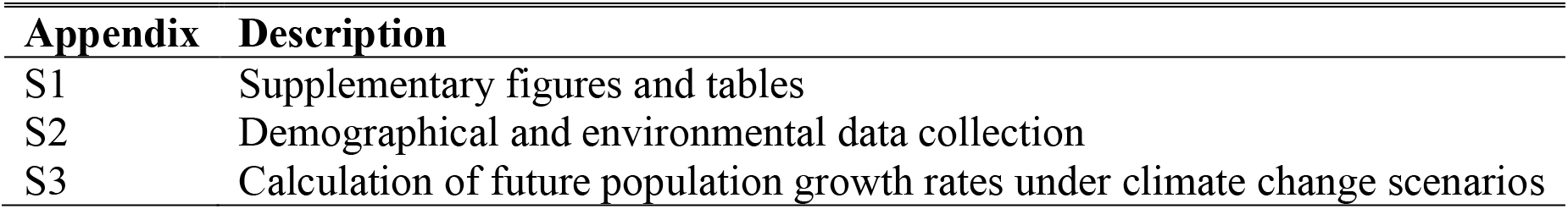

| Appendix | Description |
| --- | --- |
| S1 | Supplementary figures and tables |
| S2 | Demographical and environmental data collection |
| S3 | Calculation of future population growth rates under climate change scenarios |

### Supporting Information

Gesa Römer, Ditte M. Christiansen, Hendrik de Buhr, Kristoffer Hylander, Owen R. Jones, Sonia Merinero, Kasper Reitzel, Johan Ehrlén & Johan P. Dahlgren. 2020. Drivers of large-scale spatial demographic variation in a perennial plant. *Ecology*

## Appendix S1

**Figure S1.**
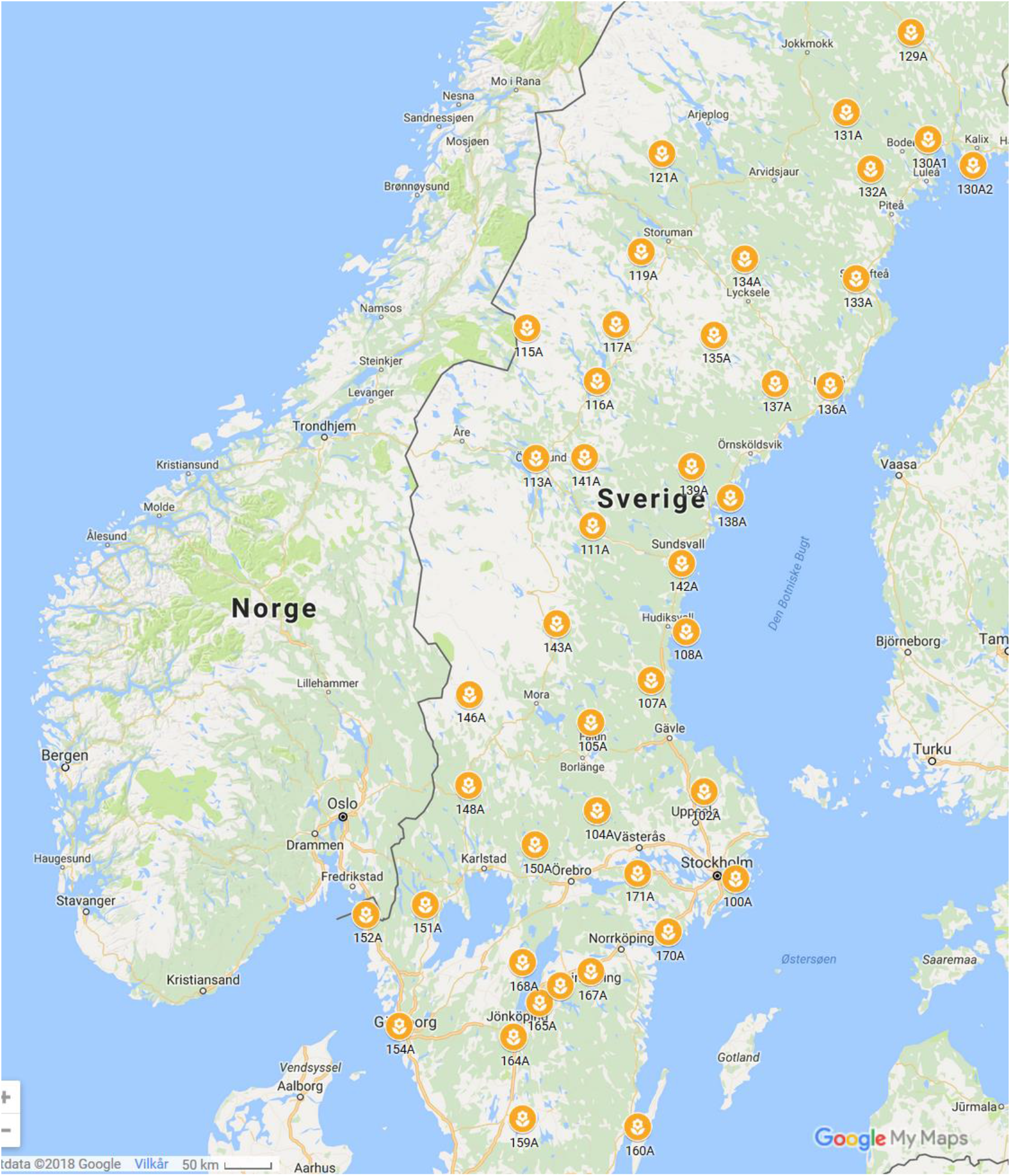
Map of all initial fieldwork sites of *Actaea spicata* in Sweden. Sites are distributed over the whole climatic range of Sweden and we tried to spread them out evenly both over latitudinal and longitudinal gradients. Site 100A was lost due to grazing in 2018, site 170A was heavily frequented by wild boar in 2018 and 2019 and Site 138A could not be accessed in 2018 and 2019. Those sites were therefore removed from the analyses.

**Figure S2:**
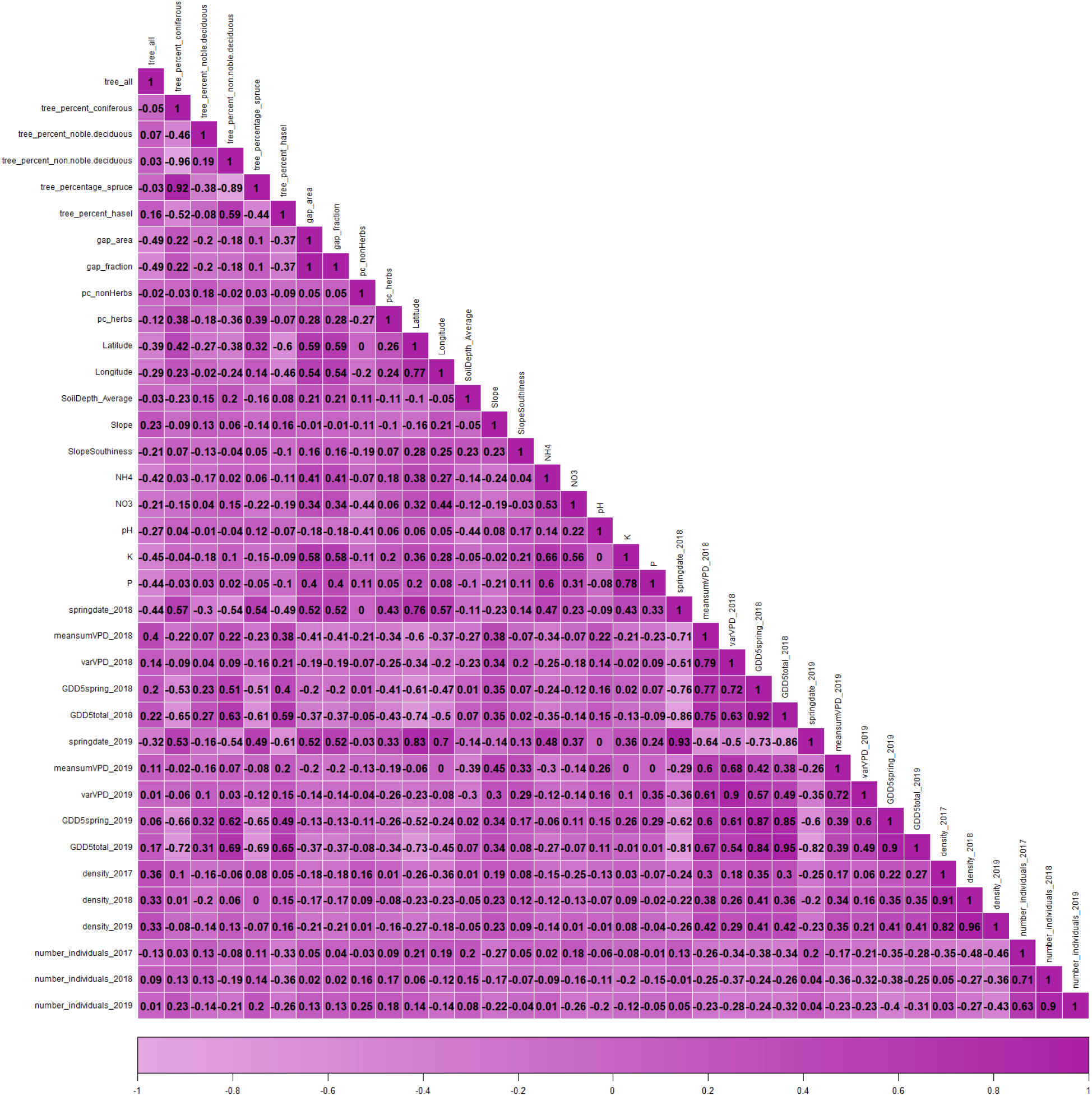
Correlation table with all environmental variables initially included in the analyses. Variables that have been collected over several years are included for each of those years and are naturally highly correlated.

**Figure S3:**
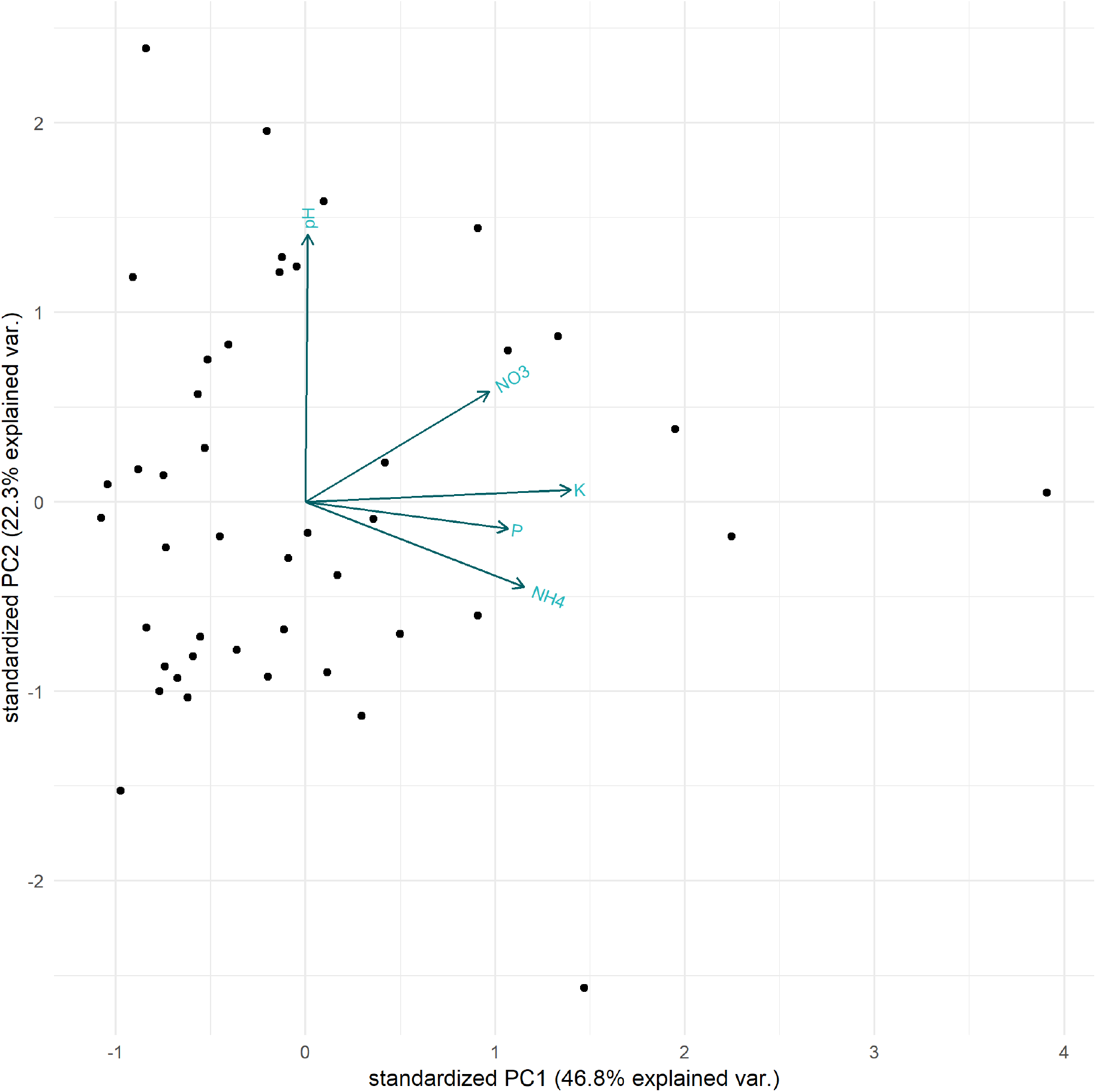
Results from Principal component analysis of the soil variables.

**Figure S4.**
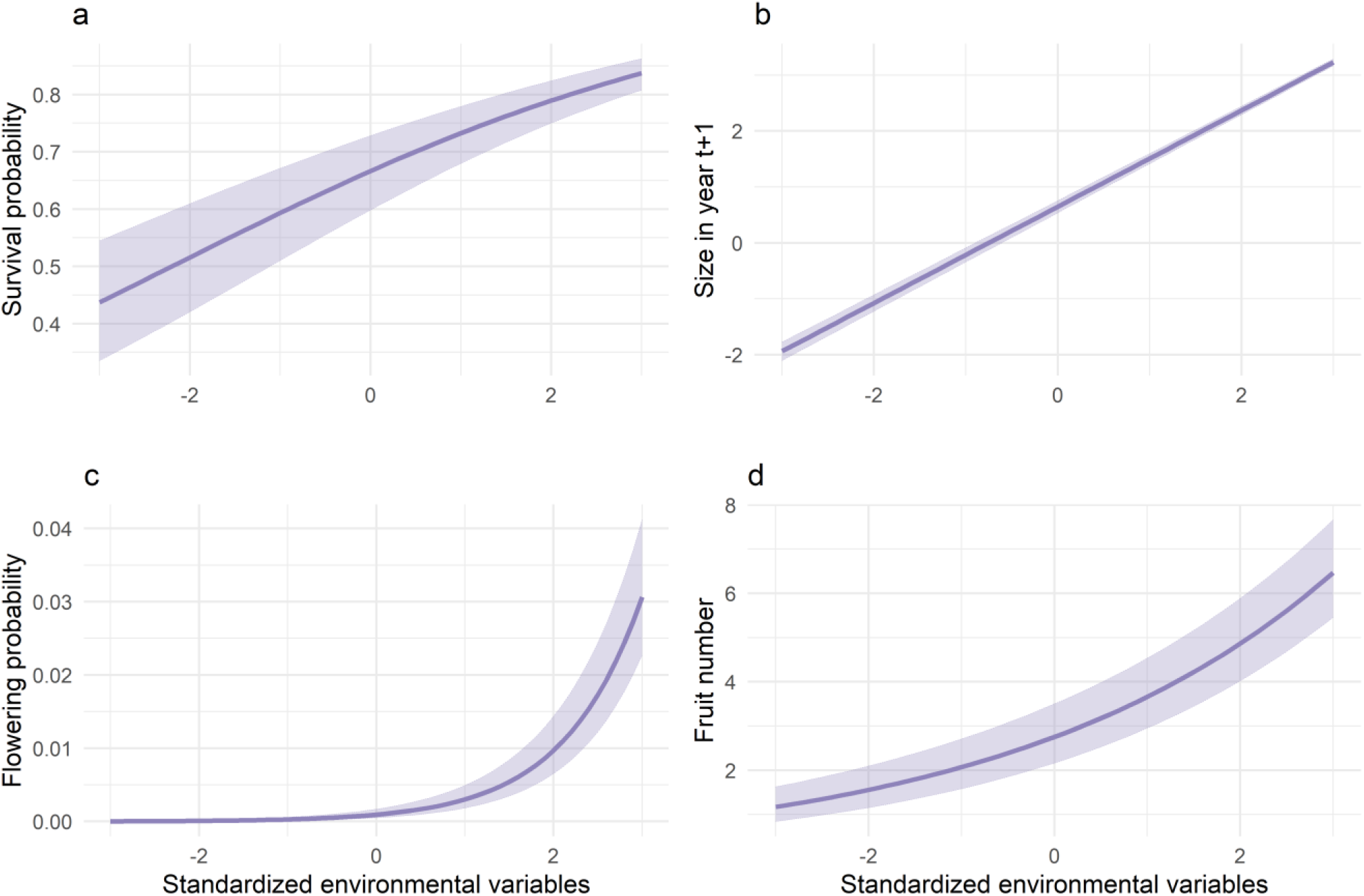
Influence of size on vital rates of A. spicata. Panel (a) shows the influence of size on survival (β = 0.31, ± = 0.03 SEM, P = < 0.01), (b) on growth (β = 0.86, ± = 0.01 SEM, t = 83.8), (c) on the probability of flowering (β = 1.17, ± = 0.05 SEM, *P* = < 0.01) and (d) on the number of fruits (β = 0.28, ± = 0.02 SEM, *P* = < 0.01). For all relationships the 95% CI is included in the figures.

**Figure S5:**
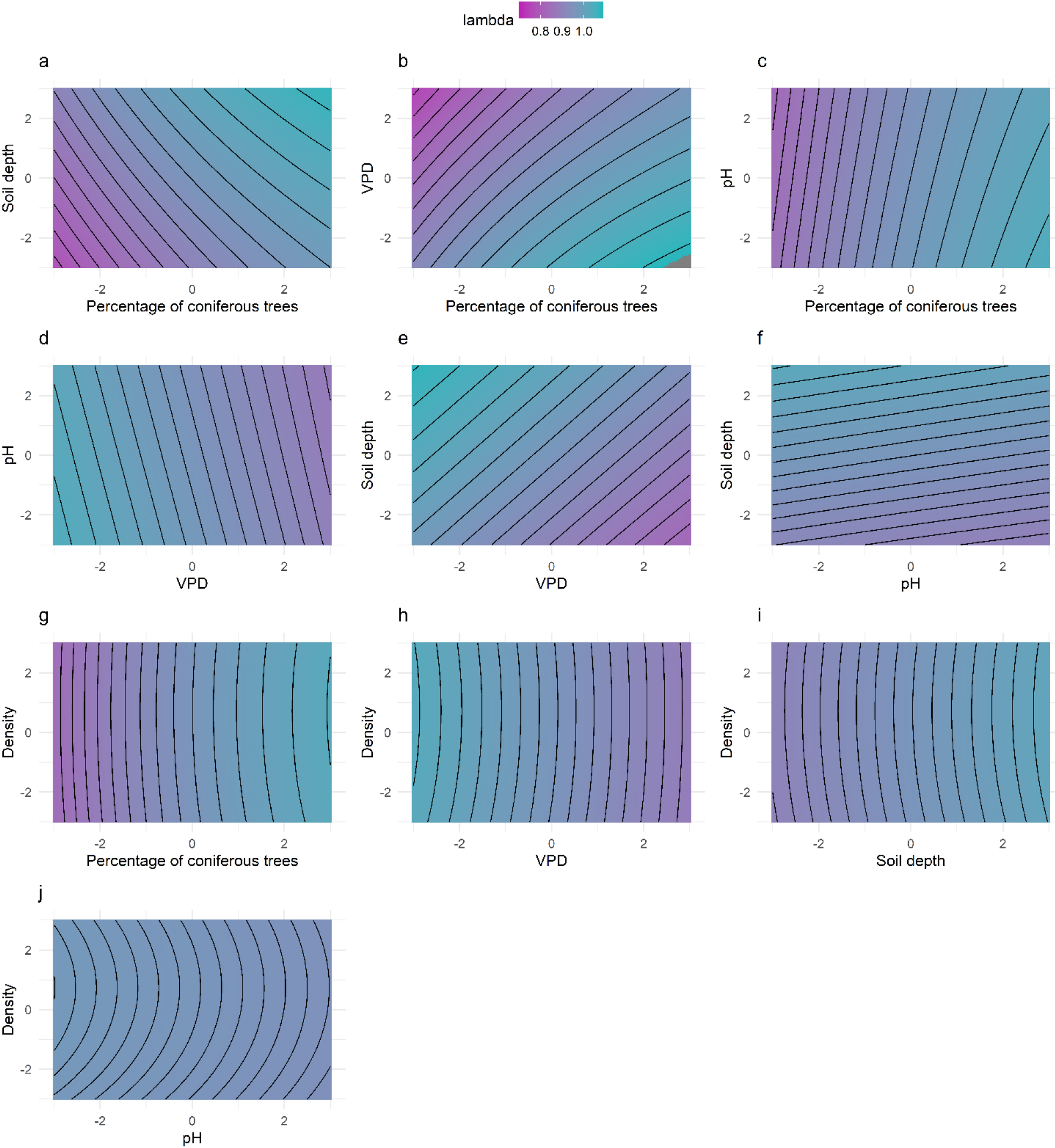
Effects of pairs of environmental variables on population growth rate. We analyzed two drivers simultaneously by evaluating *λ* for every possible combination of the range of the pair of drivers to produce a matrix of *λ*-values. The top row shows the effect of percentage of coniferous trees on the x-axis combined with (a) soil depth, (b) VPD and (c) pH. While the second row shows VPD on the x-axis combined with (d) pH and (e) soil depth. (f) Combined effect of pH and soil depth. The bottom two rows show effects of density combined with (g) percentage of coniferous trees, (h) VPD, (i) soil depth and (j) pH. All environmental variables are standardized by subtracting the arithmetic mean and dividing by the standard deviation

**Table S1:** Significance of environmental variables in the vital rate regression models over all transitions. Estimates (with standard deviations in parentheses) are from models including all tested environmental factors, but with non-significant polynomial terms omitted. • = p < 0.1, * = p < 0.05, ** = p < 0.01, *** = p < 0.001, NS = quadratic term was not significant in the model where all non-linear terms were included and has therefore not been included in the reduced model.

| Environmental driver | Survival | Growth | Flowering | Fruits |
| --- | --- | --- | --- | --- |
| Size | 0.32 (0.03) *** | 0.86 (0.01), $t = 83.79$ | 1.17 (0.05) *** | 0.28 (0.02) *** |
| VPD | -0.07 (0.11) | -0.07 (0.03), $t = -2.21$ | -0.26 (0.5) ** | -0.05 (0.04) |
| VPD<sup>2</sup> | NS | NS | NS | NS |
| K | -0.11 (0.13) | -0.001 (0.03), $t = -0.026$ | -0.29 (0.15) * | 0.07 (0.05) |
| K<sup>2</sup> | NS | NS | 0.08 (0.05) • | NS |
| pH | 0.17 (0.11) | 0.01 (0.03), $t = 0.305$ | -0.24 (0.09) ** | 0.04 (0.04) |
| Soil depth | 0.19 (0.11) • | 0.06 (0.03), $t = 2.071$ | -0.002 (0.08) | -0.006 (0.04) |
| % coniferous trees | 0.38 (0.11) *** | -0.01 (0.03), $t = -0.495$ | 0.1 (0.08) | -0.04 (0.03) |
| % coniferous trees<sup>2</sup> | NS | NS | NS | 0.15 (0.06) ** |
| Gap fraction | -0.04 (0.14) | 0.02 (0.04), $t = 0.604$ | -0.08 (0.1) | -0.002 (0.05) |
| Slope | -0.03 (0.08) | -0.002 (0.02), $t = -0.109$ | -0.05 (0.06) | -0.07 (0.04) • |
| Density | -0.01 (0.12) | 0.05 (0.03), $t = -0.333$ | -0.04 (0.09) | 0.17 (0.07) * |
| Density<sup>2</sup> | NS | NS | NS | -0.12 (0.04) ** |

**Table S2:** Population growth rates (λ) for all populations shown for all transitions pooled and for both transitions individually. Direction shows the difference between λ-values in the two individual transitions, where “+” indicates an increase in λ from the transition 2017-2018 to the transition 2018-2019 and “-” indicates a decrease. Values were calculated without including environmental variables.

| **Site** | **Population growth rate ($\lambda$)** |  |  | **direction** |
| --- | --- | --- | --- | --- |
|  | **2017-2019** | **2017-2018** | **2018-2019** |  |
| 102A | 0.84 | 0.93 | 0.85 | - |
| 104A | 0.9 | 1.06 | 0.73 | - |
| 105A | 1.05 | 1.09 | 0.98 | - |
| 107A | 0.91 | 1.03 | 0.82 | - |
| 108A | 0.99 | 1.11 | 0.84 | - |
| 111A | 1.01 | 1.09 | 0.89 | - |
| 113A | 1.04 | 1.11 | 0.95 | - |
| 115A | 0.96 | 1.04 | 0.87 | - |
| 116A | 0.90 | 1.01 | 0.84 | - |
| 117A | 0.96 | 1.03 | 0.89 | - |
| 119A | 0.94 | 1.02 | 0.83 | - |
| 121A | 1.00 | 1.06 | 0.90 | - |
| 129A | 0.97 | 1.09 | 0.83 | - |
| 130A1 | 0.87 | 0.94 | 0.81 | - |
| 130A2 | 0.95 | 1.03 | 0.80 | - |
| 131A | 0.95 | 0.94 | 0.90 | - |
| 132A | 0.94 | 0.96 | 0.90 | - |
| 133A | 0.98 | 1.03 | 0.92 | - |
| 134A | 0.96 | 1.04 | 0.92 | - |
| 135A | 0.94 | 1.02 | 0.87 | - |
| 136A | 0.96 | 1.03 | 0.83 | - |
| 137A | 0.96 | 0.99 | 0.89 | - |
| 139A | 0.97 | 1.03 | 0.88 | - |
| 141A | 0.95 | 1.03 | 0.89 | - |
| 142A | 0.88 | 1.01 | 0.88 | - |
| 143A | 0.98 | 1.08 | 0.84 | - |
| 146A | 1.00 | 1.08 | 0.90 | - |
| 148A | 0.99 | 1.03 | 0.91 | - |
| 150A | 0.95 | 0.99 | 0.83 | - |
| 151A | 0.96 | 1.09 | 0.79 | - |
| 152A | 1.02 | 1.04 | 0.88 | - |
| 154A | 0.91 | 1.03 | 0.80 | - |
| 159A | 0.96 | 1.05 | 0.87 | - |
| 160A | 0.88 | 1.00 | 0.81 | - |
| 164A | 0.99 | 1.07 | 0.91 | - |
| 165A | 0.97 | 0.96 | 0.93 | - |
| 166A | 0.95 | 1.05 | 0.83 | - |
| 167A | 0.95 | 1.04 | 0.86 | - |
| 168A | 1.02 | 1.05 | 0.93 | - |
| 171A | 1.01 | 1.09 | 0.92 | - |
| average | 0.96 | 1.04 | 0.89 | - |

**Table S3:** Overview of the influence from pairwise combinations of significant environmental drivers on population growth rates.

| Combination | Min $\lambda$ | Max $\lambda$ | Mean $\lambda$ |
| --- | --- | --- | --- |
| Density + coniferous trees | 0.84 | 1.03 | 0.95 |
| Density + VPD | 0.88 | 1.03 | 0.96 |
| Density + soil depth | 0.90 | 1.01 | 0.96 |
| Density + pH | 0.95 | 0.98 | 0.96 |
| Coniferous trees + pH | 0.83 | 1.04 | 0.95 |
| Coniferous trees + soil depth | 0.78 | 1.07 | 0.95 |
| Coniferous trees + VPD | 0.75 | 1.09 | 0.95 |
| VPD + pH | 0.88 | 1.04 | 0.96 |
| VPD + soil depth | 0.83 | 1.07 | 0.96 |
| pH + soil depth | 0.89 | 1.03 | 0.96 |

**Table S4:**
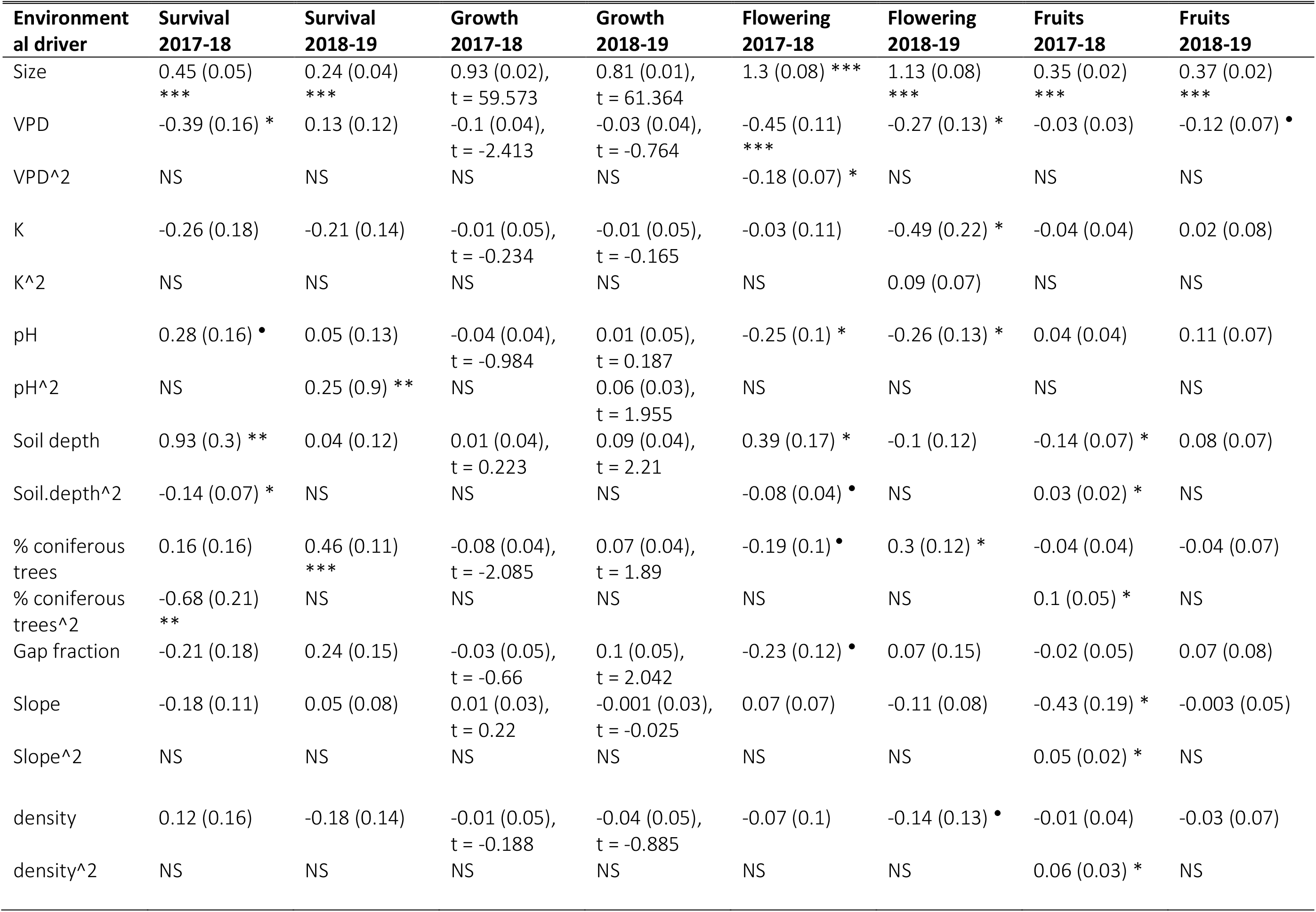
Significance of environmental variables in the vital rate regression models for the individual transitions. Estimates (with standard deviations in parentheses) are from models including all tested environmental factors, but with non-significant polynomial terms omitted. • = p < 0.1, * = p < 0.05, ** = p < 0.01, *** = p < 0.001, NS = quadratic term was not significant in the model where all non-linear terms were included and has therefore not been included in the reduced model.

| Environmental driver | Survival 2017-18 | Survival 2018-19 | Growth 2017-18 | Growth 2018-19 | Flowering 2017-18 | Flowering 2018-19 | Fruits 2017-18 | Fruits 2018-19 |
| --- | --- | --- | --- | --- | --- | --- | --- | --- |
| Size | 0.45 (0.05) *** | 0.24 (0.04) *** | 0.93 (0.02),<br>t = 59.573 | 0.81 (0.01),<br>t = 61.364 | 1.3 (0.08) *** | 1.13 (0.08) *** | 0.35 (0.02) *** | 0.37 (0.02) *** |
| VPD | -0.39 (0.16) * | 0.13 (0.12) | -0.1 (0.04),<br>t = -2.413 | -0.03 (0.04),<br>t = -0.764 | -0.45 (0.11) *** | -0.27 (0.13) * | -0.03 (0.03) | -0.12 (0.07) • |
| VPD<sup>2</sup> | NS | NS | NS | NS | -0.18 (0.07) * | NS | NS | NS |
| K | -0.26 (0.18) | -0.21 (0.14) | -0.01 (0.05),<br>t = -0.234 | -0.01 (0.05),<br>t = -0.165 | -0.03 (0.11) | -0.49 (0.22) * | -0.04 (0.04) | 0.02 (0.08) |
| K<sup>2</sup> | NS | NS | NS | NS | NS | 0.09 (0.07) | NS | NS |
| pH | 0.28 (0.16) • | 0.05 (0.13) | -0.04 (0.04),<br>t = -0.984 | 0.01 (0.05),<br>t = 0.187 | -0.25 (0.1) * | -0.26 (0.13) * | 0.04 (0.04) | 0.11 (0.07) |
| pH<sup>2</sup> | NS | 0.25 (0.9) ** | NS | 0.06 (0.03),<br>t = 1.955 | NS | NS | NS | NS |
| Soil depth | 0.93 (0.3) ** | 0.04 (0.12) | 0.01 (0.04),<br>t = 0.223 | 0.09 (0.04),<br>t = 2.21 | 0.39 (0.17) * | -0.1 (0.12) | -0.14 (0.07) * | 0.08 (0.07) |
| Soil.depth<sup>2</sup> | -0.14 (0.07) * | NS | NS | NS | -0.08 (0.04) • | NS | 0.03 (0.02) * | NS |
| % coniferous trees | 0.16 (0.16) | 0.46 (0.11) *** | -0.08 (0.04),<br>t = -2.085 | 0.07 (0.04),<br>t = 1.89 | -0.19 (0.1) • | 0.3 (0.12) * | -0.04 (0.04) | -0.04 (0.07) |
| % coniferous trees<sup>2</sup> | -0.68 (0.21) ** | NS | NS | NS | NS | NS | 0.1 (0.05) * | NS |
| Gap fraction | -0.21 (0.18) | 0.24 (0.15) | -0.03 (0.05),<br>t = -0.66 | 0.1 (0.05),<br>t = 2.042 | -0.23 (0.12) • | 0.07 (0.15) | -0.02 (0.05) | 0.07 (0.08) |
| Slope | -0.18 (0.11) | 0.05 (0.08) | 0.01 (0.03),<br>t = 0.22 | -0.001 (0.03),<br>t = -0.025 | 0.07 (0.07) | -0.11 (0.08) | -0.43 (0.19) * | -0.003 (0.05) |
| Slope<sup>2</sup> | NS | NS | NS | NS | NS | NS | 0.05 (0.02) * | NS |
| ***Actaea spicata* – demography** |  |  |  |  |  |  |  |  |
| density | 0.12 (0.16) | -0.18 (0.14) | -0.01 (0.05),<br>t = -0.188 | -0.04 (0.05),<br>t = -0.885 | -0.07 (0.1) | -0.14 (0.13) • | -0.01 (0.04) | -0.03 (0.07) |
| density^2 | NS | NS | NS | NS | NS | NS | 0.06 (0.03) * | NS |

## Appendix S2

### Section 1: Demographic data collection

#### Size data

The size of individuals of *Actaea spicata* was calculated from plant height and stem diameter following Dahlgren and Ehrlén (2009) and as given in

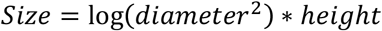

Where height is defined by the distance from the ground to the horizontal plane formed by the largest leaves. To measure this, we used a ruler starting with 0 and measure from ground to horizontal plane, ignoring the inflorescences (Fig. S1). Stem diameter was measured on the vegetative stem, 1 cm below the point, where the different leaves branch. The diameter of *A. spicata* is not round but oval. We therefore measured the point with the largest width as shown in Fig. S2.

**Figure S1:**
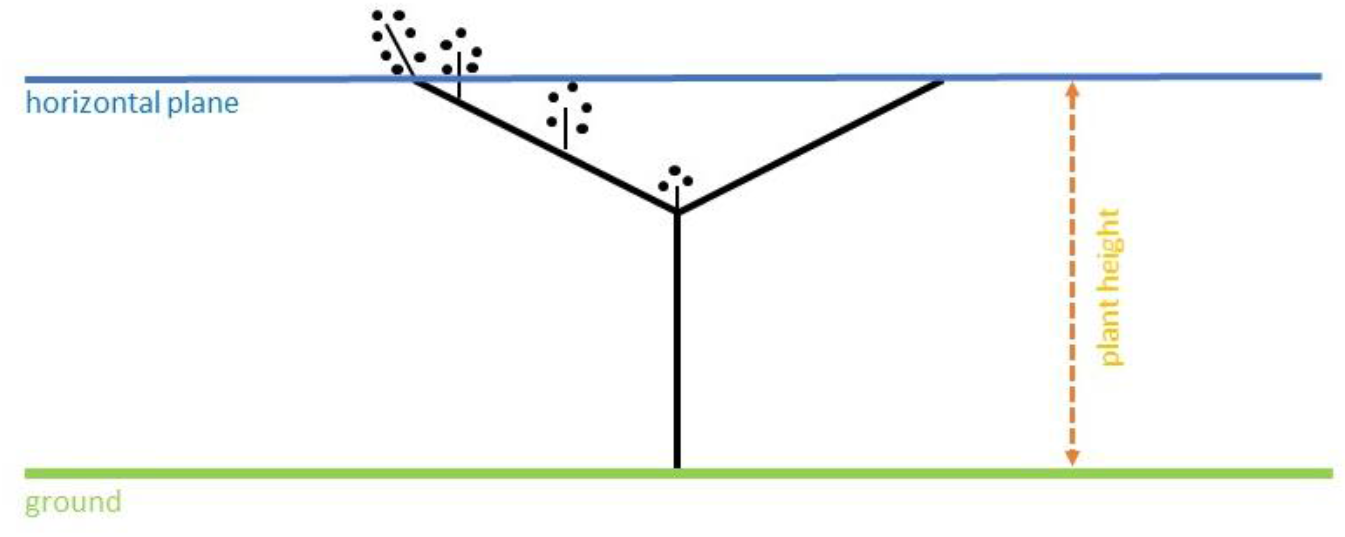
Measurement of plant height for *Actaea spicata*

**Figure S2:**
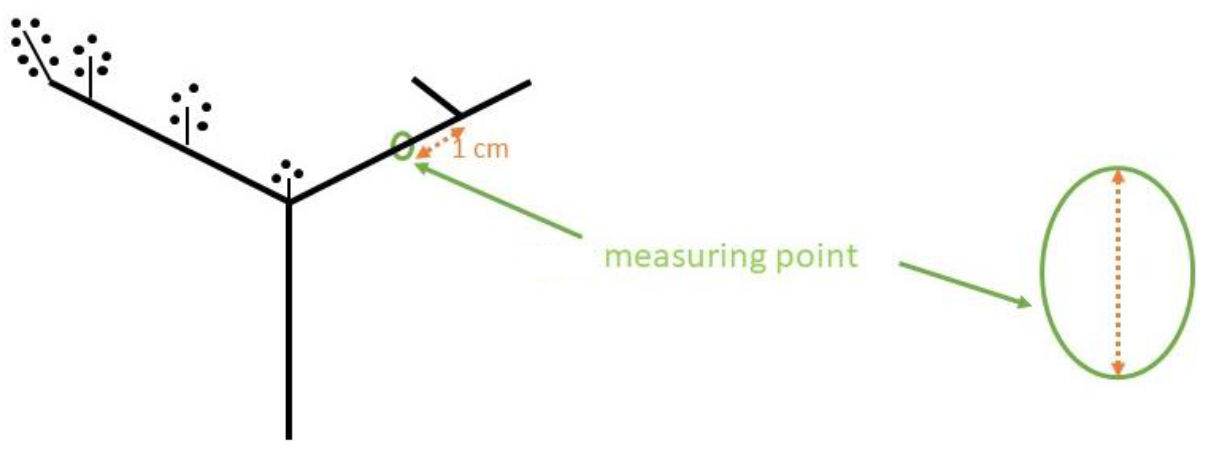
Measurement of shoot diameter for *Actaea spicata*

#### Fecundity data

*Actaea spicata* has up to four inflorescences, sometimes even more. The inflorescences do not start growing at the same time and therefore were in various stages during the census (e.g. flowering or already having fruits). The order in which inflorescences appear is shown in Fig. S3. We recorded the number of flowers or fruits in each inflorescence separately.

**Figure S3:**
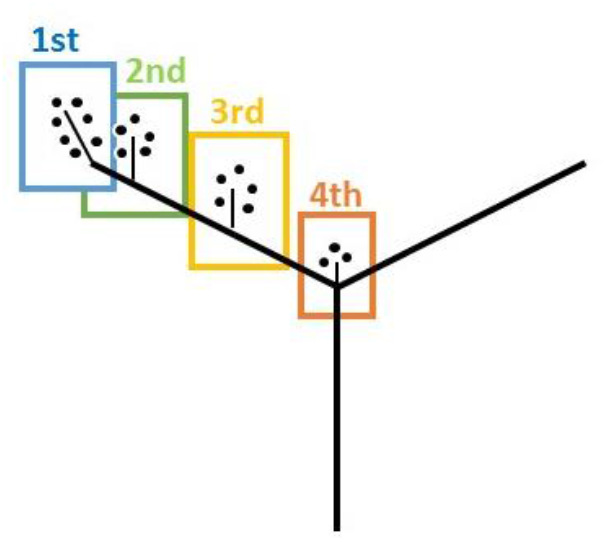
Flowering-order of Actaea spicata

### Section 2: Environmental variable collection

In addition to the demographic data we collected a range of environmental variables regarding climate, soil nutrients, topography and plant community for each site. We collected weather data throughout the whole study period. Air-temperature and humidity loggers (EasyLog EL-USB-2 from Lascar electronics) were placed in the center of each plot, tied to a wooden pole and protected from rain with a plastic mug equipped with holes to provide air circulation (as described in Terando et al. 2017). We used the logger data to calculate growing degree days with a base temperature of 5 °C for the spring period 15.05-15.06 and first spring date by the definition from the Swedish meteorological office (SMHI 2011), which is seven days in a row with mean temperatures above 0 °C after the 15th of February. In addition, we calculated Vapor Pressure Deficiency, a temperature corrected measure for moisture in the air (Jones 2014), during summer (15.06-15.08, hereafter VPD) using the R package *plantecophys* (Duursma 2015) since it is highlighted as more important for plants compared to relative humidity (Ashcroft and Gollan 2013).

In 2018 we took soil samples from five different spots within a site. The soil was taken from around 10 cm below ground (after scraping away litter from the surface). The spots were chosen so that they are spread out evenly over the site (e.g. like on a dice). If possible, the soil was taken from inside the patch. In case of a very small patch or a very dense population, we took samples from the closest possible location. Soil samples from one site were mixed and stored in a plastic-zip bag where we pressed out the air before storing them in a cooler. The samples were then analyzed in the laboratory at the University of Southern Denmark. In 2018 we collected soil samples and analyzed the concentrations of several important plant nutrients. Nitrate (NO_3_^-^) and exchangeable ammonia (NH_4_^+^) was extracted from fresh soil by a 2M KCl solution (Keeney 1982). Both NO_3_^-^ and NH_4_^+^ were analyzed spectrophotometrically on a SKALAR SAN^plus^ Analyzer. Plant available Phosphorus (P) was extracted from fresh soil by a 0.5M NaHCO_3_ solution at pH 8.5 (Olsen 1954) and phosphate (PO_4_^3-^) was determined spectrophotometrically, according to Koroleff and Grasshof (1983). Plant available potassium (K) was extracted from air dried soil by a mixture of 0.10M ammonium lactate and 0.40M acetic acid at pH 3.75 for 90 minutes (Egnér et al. 1960) and analyzed on an ICP OES (Perkin Elmer Optima 2100 DV instrument). Soil pH was measured pH on a suspension of fresh soil and water. In addition, we measured soil depth using a simple metal stick which was put into the ground towards the center of earth – disregarding potential slopes of the side. As with the soil samples, we took at least five soil depth samples, spread out like on a dice, minimizing the distance from each plant to measurement-points.

The collected topographic information including exact latitude and longitude positions of the populations, taken with a Garmin GPS map 64s when standing in the middle of the plot as well as the slope inclination of the site measured using a digital mechanic’s level in the android app *Bubble level galaxy.* Slope inclination was measured from the highest point of the site to the lowest in the way that one person stood at the highest point of the site, the other person on the lowest point. The upper person will look at a point on the lower person that matches its own eye height. Using the mechanic’s level, we can write down the approximate slope of the site (Fig. S4 and S5). Aspect direction of the slope was determined using a compass. Slope inclination and aspect direction were used to determine the variable southwards slope, calculated as standardized aspect of the site multiplied with the standardized aspect direction as distance to South, were 0 indicated a slope towards South while 190 indicates a slope aspect towards North. This variable describes the extent to which the ground was slanted towards the south and thereby acts as an additional indicator for the exposure to light and moisture. In addition, we measured site area and calculated intraspecific plant density.

In 2018 we took canopy cover pictures using the back camera of a Sony Xperia L1 with an attached fisheye-lens (180° Supreme Fisheye Lens, Model MFE4 by MPOW Inc). Pictures were taken pointing away from the earth center perpendicular to the ground, disregarding the potential slope of the site, from 60 cm above ground using a stick to ensure the same distance for every picture. Those images were then processed in ImageJ (Schneider et al. 2012) using the plugin Hemispherical 2.0 (Beckschäfer 2015) to calculate the gap fraction. If different parts of the site had substantially different canopy covers, several pictures from different spots within the site were taken and then averaged.

Lastly, we collected data on the plant community. We defined tree species and their abundance using a relascope, a forestry instrument to measure the basal area, for each tree visible from the center of the site to determine the percentage of coniferous vs. broadleaf trees. Looking through the relascope, a tree that is smaller than the relascope opening was counted, one that exactly fits the opening counted as 0.5, and trees bigger than the opening are counted as one (Fig. S6). Each stem was counted individually at the height of the relascope (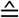 eye height). One exception for this procedure were hazel trees since they often grow in a cluster with stems being too small to be measured in the traditional way. Instead, we measured the stem diameter, as we expected it to be if all stems were tightly layered together. Trees that were smaller than 5 cm in diameter were not counted.

**Figure S4:**
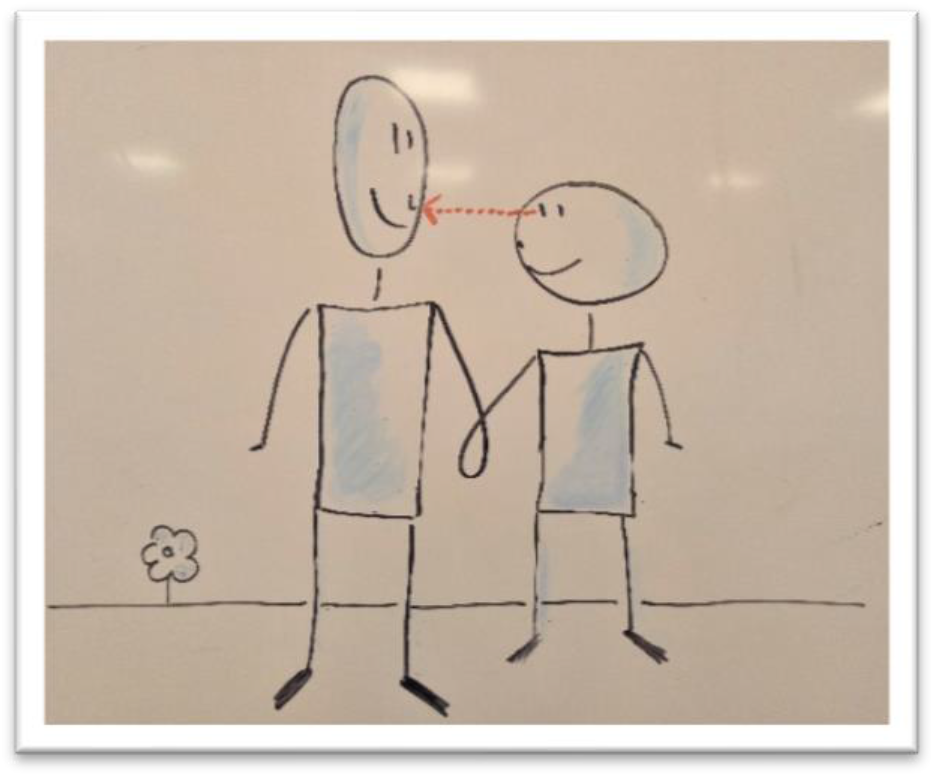
Upper and Lower are standing on even ground.

**Figure S5:**
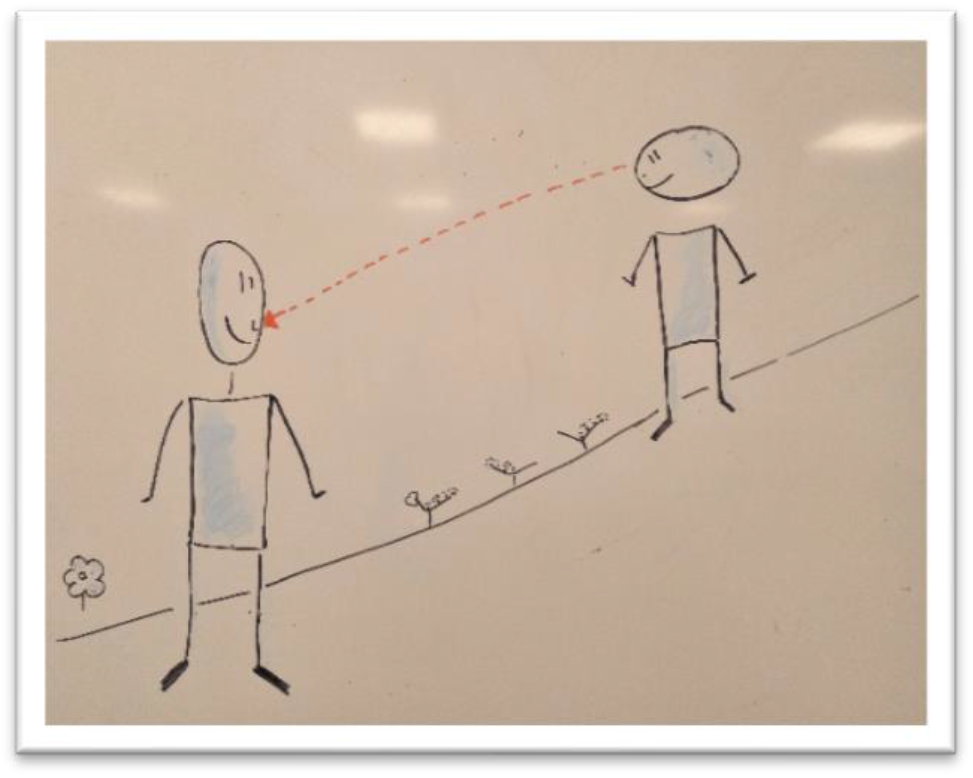
Upper stands at the highest point, Lower at the lowest.

**Figure S6:**
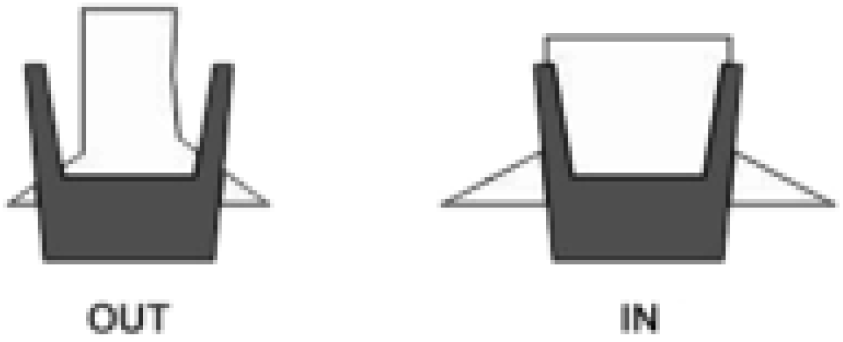
Looking through a relascope the left tree would not count; the right tree would be counted as 0.5; trees bigger are counted as 1.

## Appendix S3 Calculation of future population growth rates under climate change scenarios

To get an estimate of how population growth rates (*λ*) of *Actaea spicata* will change in the future, we used temperature estimates for Sweden from a report of the Swedish meteorological office (Sjökvist et al. 2015). The report uses the IPCC scenario RCP 4.5 (IPCC 2014) as baseline for their calculations and predicted an increase in temperature of 2°C in southern Sweden and 4°C in northern Sweden until the end of the century (Sjökvist et al. 2015). For our calculations we used the mean prognosed temperature increase of 3°C. As an estimate of the changes in humidity we used a report about changes in air-humidity stating that relative humidity increased by roughly 1% in the past 50 years (Wern 2013). We then calculated a prognosed vapor pressure deficit (VPD) value for the year 2100 from those estimates and repeated our analyses with this value, leading to an increased average population growth rates of *λ* = 1.01 (compared to *λ* = 0.96 during our study period).

## Literature Cited

Anderberg A., and Anderberg A.-L. 2017. Den virtuella floran. Naturhistoriska riksmuseet. http://linnaeus.nrm.se/flora/ (February 03, 2020).

Anderson, D. B. 1936. Relative Humidity or Vapor Pressure Deficit. Ecology 17:277–282.

Araújo, M. B., and C. Rahbek. 2006. Ecology. How does climate change affect biodiversity? Science (New York, N.Y.) 313:1396–1397.

Bates, D., M. Mächler, B. Bolker, and S. Walker. 2015. Fitting Linear Mixed-Effects Models Using lme4. Journal of Statistical Software 67.

Beckschäfer, P. 2015. Hemispherical_2.0 – Batch processing hemispherical and canopy photographs with ImageJ – User Manual. Chair of Forest Inventory and Remote Sensing, Georg-August-Universität Göttingen, Germany.

Bruna, E. M., and M. K. Oli. 2005. Demographic effects of habitat fragmentation on a tropical herb: Life-Table Response Experiments. Ecology 86:1816–1824.

Chave, J. 2013. The problem of pattern and scale in ecology: what have we learned in 20 years? Ecology letters 16:4–16.

Csergő, A. M., R. Salguero-Gómez, O. Broennimann, S. R. Coutts, A. Guisan, A. L. Angert, E. Welk, I. Stott, B. J. Enquist, B. McGill, J.-C. Svenning, C. Violle, and Y. M. Buckley. 2017. Less favourable climates constrain demographic strategies in plants. Ecology letters 20:969–980.

Dahlgren, J. P., and J. Ehrlén. 2009. Linking environmental variation to population dynamics of a forest herb. Journal of Ecology 97:666–674.

Dahlgren, J. P., and J. Ehrlén. 2011. Incorporating environmental change over succession in an integral projection model of population dynamics of a forest herb. Oikos 120:1183–1190.

Diez, J. M., I. Giladi, R. Warren, and H. R. Pulliam. 2014. Probabilistic and spatially variable niches inferred from demography. Journal of Ecology 102:544–554.

Doak, D. F., and W. F. Morris. 2010. Demographic compensation and tipping points in climate-induced range shifts. Nature 467:959–962.

Duursma, R. A. 2015. Plantecophys--An R Package for Analysing and Modelling Leaf Gas Exchange Data. PloS one 10:e0143346.

Easterling, M. R., S. P. Ellner, and P. M. Dixon. 2000. Size-specific sensitivity: Applying a new structured population model. Ecology 81:694–708.

Ehrlén, J., and O. Eriksson. 2000. Dispersal limitation and patch occupancy in forest herbs. Ecology 81:1667–1674.

Ehrlén, J., and W. F. Morris. 2015. Predicting changes in the distribution and abundance of species under environmental change. Ecology letters 18:303–314.

Ehrlén, J., W. F. Morris, T. von Euler, and J. P. Dahlgren. 2016. Advancing environmentally explicit structured population models of plants. Journal of Ecology 104:292–305.

Elith, J., and J. R. Leathwick. 2009. Species Distribution Models: Ecological Explanation and Prediction Across Space and Time. Annual Review of Ecology, Evolution, and Systematics 40:677–697.

Ellner, S. P., D. Z. Childs, and M. Rees. 2016. Data-driven Modelling of Structured Populations. Springer International Publishing, Cham.

Ellner, S. P., and M. Rees. 2006. Integral projection models for species with complex demography. The American Naturalist 167:410–428.

Eriksson, O. 1995. Asynchronous flowering reduces seed predation in the perennial forest herb Actaea spicata. Oecologia 16:195–203.

Fox, J., and S. Weisberg. 2018. Visualizing Fit and Lack of Fit in Complex Regression Models with Predictor Effect Plots and Partial Residuals. Journal of Statistical Software 87.

Fröborg, H., and O. Eriksson. 2003. Predispersal seed predation and population dynamics in the perennial understorey herb Actaea spicata. Canadian Journal of Botany 81:1058–1069.

Garten, C. f., W. M. Post, P. J. Hanson, and L. W. Cooper. 1999. Forest soil carbon inventories and dynamics along an elevation gradient in the southern Appalachian Mountains. Biogeochemistry 45:115–145.

Givnish, T. J. 1987. Comparative studies of leaf form: assessing the relative roles of selection pressures and phylogenetic constraints. New Phytologist 106:131–160.

Greiser, C., K. Hylander, E. Meineri, M. Luoto, and J. Ehrlén. 2020. Climate limitation at the cold edge: contrasting perspectives from species distribution modelling and a transplant experiment. Ecography 11:36.

Guisan, A., and W. Thuiller. 2005. Predicting species distribution: offering more than simple habitat models. Ecology Letters 8:993–1009.

Harrell, F. E. 2001. Overfitting and limits on number of predictors. Regression Modeling Strategies:60–64.

Jones, H. G. 2014. Plants and microclimate. A quantitative approach to environmental plant physiology. Cambridge University Press, Cambridge, New York.

Levin, S. A. 1992. The Problem of Pattern and Scale in Ecology: The Robert H. MacArthur Award Lecture. Ecology 73:1943–1967.

Lükewille, A., and R. Wright. 1997. Experimentally increased soil temperature causes release of nitrogen at a boreal forest catchment in southern Norway. Global Change Biology 3:13–21.

Merow, C., S. T. Bois, J. M. Allen, Y. Xie, and J. A. Silander. 2017. Climate change both facilitates and inhibits invasive plant ranges in New England. Proceedings of the National Academy of Sciences of the United States of America 114:E3276–E3284.

Merow, C., J. P. Dahlgren, C. J. E. Metcalf, D. Z. Childs, M. E.K. Evans, E. Jongejans, S. Record, M. Rees, R. Salguero-Gómez, and S. M. McMahon. 2014a. Advancing population ecology with integral projection models: a practical guide. Methods in Ecology and Evolution 5:99–110.

Merow, C., A. M. Latimer, A. M. Wilson, S. M. McMahon, A. G. Rebelo, and J. A. Silander. 2014b. On using integral projection models to generate demographically driven predictions of species’ distributions: development and validation using sparse data. Ecography 37:1167–1183.

Morris, W. F., J. Ehrlén, J. P. Dahlgren, A. K. Loomis, and A. M. Louthan. 2020. Biotic and anthropogenic forces rival climatic/abiotic factors in determining global plant population growth and fitness. Proceedings of the National Academy of Sciences of the United States of America 117:1107–1112.

Mossberg, B., and L. Stenberg. 2014. Den nye nordiske flora. Gyldendal, Kbh.

Nicolè, F., J. P. Dahlgren, A. Vivat, I. Till-Bottraud, and J. Ehrlén. 2011. Interdependent effects of habitat quality and climate on population growth of an endangered plant. Journal of Ecology 99:1211–1218.

Pellmyr, O. 1984. The pollination ecology of Actaea spicata (Ranunculaceae). Nordic Journal of Botany 4:443–456.

Pfister, C. A. 1998. Patterns of variance in stage-structured populations: evolutionary predictions and ecological implications. Proceedings of the National Academy of Sciences 95:213–218.

R Core Team. 2018. R: A Language and Environment for Statistical Computing. R Foundation for Statistical Computing. https://www.R-project.org.

Rees, M., D. Z. Childs, J. C. Metcalf, K. E. Rose, A. W. Sheppard, and P. J. Grubb. 2006. Seed dormancy and delayed flowering in monocarpic plants: selective interactions in a stochastic environment. The American Naturalist 168:E53–E71.

Schneider, C. A., W. S. Rasband, and K. W. Eliceiri. 2012. NIH Image to ImageJ: 25 years of image analysis. Nature methods 9:671–675.

Silvertown, J., M. Franco, I. Pisanty, and A. Mendoza. 1993. Comparative Plant Demography--Relative Importance of Life-Cycle Components to the Finite Rate of Increase in Woody and Herbaceous Perennials. Journal of Ecology 81:465–476.

Sjökvist E., J. Axén Mårtensson, J. Dahné, N. Köplin, E. Björck, L. Nylén, J. Tengdelius Brunell, D. Nordborg, K. Hallberg, and J. Södling. 2015. Klimatscenarier för Sverige: Bearbetning av RCP-scenarier för meteorologiska och hydrologiska effektstudier. SMHI.

Sletvold, N., J. P. Dahlgren, D.-I. Oien, A. Moen, and J. Ehrlén. 2013. Climate warming alters effects of management on population viability of threatened species: results from a 30-year experimental study on a rare orchid. Global Change Biology 19:2729–2738.

SMHI. 2011. Vår. SMHI. https://www.smhi.se/kunskapsbanken/meteorologi/var-1.1080 (February 03, 2020).

Wern, L. 2013. Luftfuktighet. Variationer i Sverige.

Zeipel H. von, O. Eriksson, and J. Ehrlén. 2006. Host plant population size determines cascading effects in a plant–herbivore–parasitoid system. Basic and Applied Ecology 7:191–200.

## Literature Cited

Ashcroft, M. B., and J. R. Gollan. 2013. Moisture, thermal inertia, and the spatial distributions of nearsurface soil and air temperatures: Understanding factors that promote microrefugia. Agricultural and Forest Meteorology 176:77–89.

Egnér, H., H. Riehm, and W. R. Domingo. 1960. Investigations on chemical soil analysis as the basis for estimating soil fertility. II. Chemical extraction methods for phosphorus and potassium determination. Kungliga Lantbrukshögskolans Annaler 26:199–215.

Keeney. 1982. Nitrogen-Inorganic form. In Methods of soil analysis (part 2) Chemical and mirobiological properties. American Society of Agronomy:1159, 643–698.

Koroleff, F., and K. Grasshof. 1983. Determination of nutrients, 125–188. Methods of seawater analyses, edited by: Grasshof, K., Erhardt, M., and Kremling, K., Verlag Chemie, Weinheim.

Olsen, S. R. 1954. Estimation of available phosphorus in soils by extraction with sodium bicarbonate. US Department of Agriculture.

Terando, A. J., E. Youngsteadt, E. K. Meineke, and S. G. Prado. 2017. Ad hoc instrumentation methods in ecological studies produce highly biased temperature measurements. Ecology and evolution 7:9890–9904.

## Literature Cited

IPCC. 2014. Climate change 2013. The physical science basis Working Group I contribution to the Fifth assessment report of the Intergovernmental Panel on Climate Change. Cambridge University Press, New York.

Wern, L. 2013. Luftfuktighet. Variationer i Sverige.

